# Fat body lipogenic capacity in honey bee workers is affected by age, social role, and dietary protein

**DOI:** 10.1101/2024.03.24.586478

**Authors:** Sebastian Scofield, Gro. V. Amdam

**Affiliations:** School of Life Sciences, Arizona State University, Tempe, AZ 85287, USA; Faculty of Environmental Sciences and Natural Resource Management, Norwegian University of Life Sciences, Aas, Norway

**Keywords:** bee, lipogenesis, metabolism, fatty acid, nutrient regulation, eusociality, social insects, protein, lipid

## Abstract

All organisms need to balance processes that consume energy against those that produce energy. With an increase in biological complexity over evolutionary time, regulation of this balance has become much more complex, resulting in specialization of metabolic tasks between organelles, cells, organs, and in the case of eusocial organisms, between the individuals who comprise the ‘superorganism.’ Exemplifying this, nurse honey bees maintain high abdominal lipids, while foragers have very low lipid stores, likely contributing to efficient performance of their social role, and thus to colony fitness. The proximate mechanisms responsible for these metabolic differences remain poorly understood. Here, we investigated the effects of age, age class, and dietary macronutrients on the abdominal activity of fatty acid synthase (FAS), the enzyme responsible for *de novo* synthesis of fatty acids. We found that FAS activity declines as bees age past peak nursing age. Feeding both nurses and foragers carbohydrates increased FAS activity compared with starved bee, but, whether fed or starved, nurses had much higher FAS activity than similarly treated foragers, implicating reduced lipid synthesis as one component of foragers’ low lipid stores. Finally, we used artificial diets with different amounts of protein and fat to precociously induce low, forager-like FAS activity levels in nurse-age bees deprived of protein. We speculate that reduced protein appetite and consumption during the nurse-forager transition is responsible for suppressed lipid synthesis in foragers.

**Summary statement:** Honey bee workers show reduced fat synthesis capacity as they age and leave the nest to forage. Young bees deprived of protein have low, forager-like fat synthesis capacity.

## Introduction

Metabolic homeostasis is essential for life. All organisms need to balance reactions that consume energy (anabolism) with those that produce energy (catabolism), especially in the face of temporally variable energy availability in the environment. When energy is abundant, cells generally store energy in the form of fatty acids (Renne and Hariri, 2021). These fatty acids can be taken up from the extracellular environment or synthesized in *de novo* lipogenesis by fatty acid synthase (FAS), an enzyme complex that is conserved across almost all organisms. The fatty acids are then typically stored as neutral lipids such as triacylglycerol (TAG) and sequestered in storage organelles called lipid droplets present from bacteria to metazoans. TAG represents the most concentrated form of chemically bound energy in cells and is the quantitatively most important energy reservoir in many organisms. In times of energetic stress, this energy can be released by lipolytic enzymes called lipases that break TAG molecules down into to FAs, which then become available for energy production or as building blocks for structural membrane lipids or signaling molecules (Heier and Kühnlein, 2018). The balance between energy storage by lipogenesis and energy release by lipolysis is thus of critical importance to organismal survival and is dynamically regulated in response to energetic availability and demand.

This homeostatic regulation is relatively simple in protozoans, where energy balance is regulated at a cellular level, but has become increasingly complex with the evolution of higher levels of biological organization (Lempradl, Pospisilik and Penninger, 2015), and can occur at the level of the cell, tissue, organ, and multicellular organism as a whole (Torday, 2015). Regulation at higher levels has resulted in the evolution of metabolic division of labor, which allows for the decoupling of anabolism and catabolism (Negroni and LeBoeuf, 2023). For example, the evolution of organ systems in metazoans has required development of complex regulatory systems to coordinate energy demand between different organs with specialized function (Zhao and Karpac, 2020), resulting in some tissues that specialize in lipid synthesis and storage and others that use these lipids as fuel. Some tissues thus incur a metabolic cost to provide for the energetic needs of other tissues, which can only be explained by fitness benefits at a level of organization above tissues (the individual organism) (Negroni and LeBoeuf, 2023).

Evolution of levels of organization above the individual organism is exemplified by eusociality (Szathmáry and Smith, 1995). Eusociality is a system of individuals living together in colonies and exhibiting cooperative brood care, overlap of generations, and reproductive division of labor. This division of labor typically imply that most individuals within a colony forego direct reproductive output in favor of indirect contribution to the reproduction of close relatives, such as care for the offspring of nestmates (Crespi and Yanega, 1995). Critically, because of this reproductive division of labor, the colony can become the unit of selection with fitness determined by colony-level traits (Friedman, Johnson and Linksvayer, 2020). As a result, the principles of colony physiology are broadly the same as organismal physiology, and anabolic and catabolic metabolic tasks may be distributed among individuals in a colony to increase colony-level fitness (Friedman, Johnson and Linksvayer, 2020; Negroni and LeBoeuf, 2023).

A well-studied example is found in the division of labor between queens and female workers in the Western honey bee, *Apis mellifera*. Queens, who are responsible for most or all of the reproductive output of the colony, develop in response to high nutrition (quality and amount) during the larval period (Patel *et al*., 2007), and display adult physiology consistent with high nutritional status, such as abundant lipid stores (Strachecka *et al*., 2021). The female workers, who are typically functionally sterile, develop in response to less nutrition (quality and amount) during the larval period and have lower adult lipid stores. Reproductive division of labor thus has as its basis the net transfer of energy from workers to queens, with individual workers incurring an energetic cost in order to increase not their own fitness, but the collective fitness of the colony (Negroni and LeBoeuf, 2023). This system is argued to be a key driver of the evolutionary success of eusocial species (Bernadou, Kramer and Korb, 2021).

Because they forgo direct reproduction, workers instead contribute to colony fitness through the performance of social tasks. Honey bees evolved a system of temporal polyethism, in which behavior correlates with chronological age: in the first phase of life, workers perform a series of (inside) nest tasks, most notably nursing, in which bees produce proteinaceous jelly in highly developed glands on the head to feed to developing bees and other members of the colony. Such tasks that are performed inside the protected nest carry low extrinsic mortality risk. Next, in response to colony needs and their own physiological status, workers transition from inside tasks to tasks that involve outside activities such as foraging. Foraging involves making frequent flights to collect food and water resources to nourish the colony (Robinson, 1992). Flight activity is associated with high extrinsic mortality risk (Prado *et al*., 2020).

The behavioral transition from nest tasks to foraging is accompanied by many physiological changes in workers. Mechanistically, one of these events occurs in the fat body, the major site of lipid synthesis and storage in insects. Nurses maintain high lipid stores in the fat body, but undergo dramatic loss of lipid stores prior to the transition to foraging behavior and remain lean for the remainder of their lives (Toth and Robinson, 2005). Loss of lipids is thought to be functionally important, minimizing energetic loss to the environment from the high attrition rate of foragers (Prado *et al*., 2020) and increasing flight performance by lowering body weight (Vance *et al*., 2009). The changed physiology also makes evolutionarily sense, as foragers are stripped of transferable resources that (otherwise) would be lost to the colony when foragers die in the field (Amdam and Page, 2005), increasing colony-level fitness.

Studies have shown that this lipid loss is accompanied by broad transcriptional (Ament *et al*., 2011) and proteomic changes (Chan *et al*., 2011) in the fat body but specific metabolic mechanisms remain unclear. There are a number of possibilities: lipids could be metabolically consumed to fuel the high flight activity of foragers, but this is unlikely because lipids are already low on the first day of foraging and do not correlate with foraging activity metrics (Toth and Robinson, 2005). Lower intake of dietary lipids could be important, since foragers consume much less pollen than nurses, and pollen is the bees’ primary source of dietary lipids (Crailsheim *et al*., 1992). However, evidence indicates that *de novo* lipogenesis contributes substantially to lipid stores when dietary lipids are deficient (Toth *et al*., 2005), and this does not explain why foragers, who maintain higher hemolymph sugar levels than nurses (Mayack *et al*., 2019), do not convert these sugars into lipid stores, especially since high sugar diets and resulting high hemolymph sugar levels are known to promote lipid storage in related insects such as *Drosophila* (Baenas and Wagner, 2022).

Based on these previous findings, we hypothesized that worker honey bees experience a developmental shift in the balance between lipid synthesis and breakdown during the nurse-forager transition. To test for the presence of this shift, we looked for changes in the fat body activity levels of two key enzymes, FAS and lipase, during a time frame around the transition from in-nest tasks to foraging. We then characterized the relevant contributions of feeding status and age class by starving and feeding nurses and foragers. Finally, to test whether a difference in macronutrient consumption could account for the differences we found in FAS activity, we used artificial diets to manipulate the availability of protein and lipid to workers during the first 8 days of adult life.

## Materials and methods

### Animals

Frames of sealed brood were collected from three different honey bee (*Apis mellifera ligustica*) colonies kept at Arizona State University (Tempe, AZ, USA). Brood frames were kept overnight in ventilated mesh boxes in an incubator kept at 33°C and humidified by placing an open container of water at the bottom of the incubator. Newly emerged workers were brushed from the three frames the next day and combined to form a mixed population to increase genetic diversity of the experimental groups. Bees were then used in subsequent experiments.

### Age-based sampling from colonies

To determine the effect of age on lipid metabolism, newly emerged bees were marked and released into colonies, and collected at 3, 8, and 13 days of age. On the day of sampling, workers were anaesthetized on ice, euthanized, and dissected. Bees of all age groups were collected systematically on a sampling day, and the experiment was replicated twice in two different host colonies on two different sampling days.

### Starvation of nurses and foragers collected from colonies

To determine the effect of feeding status on the lipogenic capacity of bees in two different age classes, nurses (8-day old bees observed inserting their heads into larval cells for at least 3 seconds) and foragers (28-day old bees collected when returning to the colony) were placed into (5 x 12 x 16 cm) Plexiglass and mesh cages (30 bees/cage) and fed 30% sucrose solution in 12 mL syringes *ad libitum* for 3 hours. Cages were then either fed 30% sucrose solution for another 21 hours (feeding treatment) or instead fed only water for the same period (starvation), for a total of 24 hours of caging prior to sampling. The experiment was replicated four times.

### Dietary protein and lipid deprivation of bees raised in cages

To test whether a difference in macronutrient consumption between nurses and foragers could cause the differences in FAS activity we measured in the previous experiment, we designed an experiment to measure the effect of dietary protein and lipid on lipogenic capacity in workers. Because feeding foragers nurse-like, high protein diets results in extremely high mortality (up to 100% mortality at 7 days of feeding) (Paoli *et al*., 2014), we chose instead to feed artificial diets with variable protein and lipid content to newly emerged bees until they were 8 days old, considered to be near peak nursing age. Such an approach of manipulating nurse-age bees to look for factors that precociously create forager-like physiology is commonly used to study the regulation of the nurse-forager transition (Toth *et al*., 2005; Antonio *et al*., 2008; Traniello *et al*., 2020). Bees were maintained in a humidified incubator at 31°C in cup cages (Evans *et al*., 2009). Similar to previous work (Arien, Dag and Shafir, 2018), we created diets using honey as a carbohydrate source, soy as a protein source, and corn oil as a lipid source. The composition of the diets was 70.26-78.26% honey, which contains negligible protein and lipids; 0 or 21.74% soy protein isolate (composed of 92% protein, amounting to 20% net protein in the diet); 0 or 21.74% powdered sucrose, as a replacement for the soy protein isolate in protein-deficient diets; and 0 or 8% corn oil. These percentages of dietary protein and lipid were chosen based on past work showing that these values produced bees with the highest brood-rearing capacity (Arien *et al*., 2020). A two-factor design was used for diets to test the effect of presence/absence of these concentrations of protein and lipid. The dietary mixes were fed *ad libitum* to the bees in 1.5 mL Eppendorf tubes and replaced every 24 hours. In addition, bees were fed 30% sucrose solution *ad libitum* in 10 mL syringes. Mortality was monitored daily in each cage and dead bees were removed. Consumption of the paste diets was calculated by mass changes every 24 hours. Sucrose solution consumption was measured by mass difference between the start and end of the experiment along with a correction for evaporation of the solution. All consumption metrics were calculated by dividing cage-level mass changes by the number of bees still alive at each time point. Total calories consumed by each bee per day was estimated by converting macronutrient consumption in mg/bee/day into calories, using 4, 4, and 9 calories/mg as estimations for protein, carbohydrate, and lipid, respectively. The experiment was replicated three times.

### Fat body FAS activity, lipase activity, and protein quantity in pooled samples

Because recent findings indicate that FAS is regulated post-transcriptionally by autoacetylation based on nutrient availability (Miao *et al*., 2022), we chose to measure the activity level of fat body FAS activity rather than gene expression. Fat body FAS activity was assayed using a previously-described assay (Lu, Chuang and Hsu, 2017) with minor modifications. In brief, workers were dissected and fat bodies (complete abdominal cuticle with adhering fat body tissue minus the stinger, gut, crop, and ovaries) from two bees were pooled and homogenized in 250 µL of phosphate-buffered saline (7 mM NaCl, 2.7 mM KCl, and 10 mM PO4, pH 7.4) containing cOmplete protease inhibitors (Roche) at kept on ice until further processing. The fresh samples were sonicated for 30s (Qsonica Q800R2) and centrifuged at 5000g for 10 min at 4°C. The supernatant was then collected and aliquoted: 133.2 μL was used immediately for the FAS activity assay, while the remainder was frozen and maintained at -20°C until it was used for the lipase activity and protein quantity assays.

To measure FAS activity, 33.3 μL of the supernatant was added to a buffer solution containing 163.3 μL of 2.0 M potassium phosphate buffer, pH 7.1, 16.7 μL of 20 mM dithiothreitol, 20 μL of 0.25 mM acetyl-CoA, 16.7 μL of 60 mM EDTA, and 16.7 μL of 6 mM nicotinamide adenine dinucleotide phosphate (NADPH) in a 96-well microplate. To initiate the reaction, 33.3 μL of 0.39 mM malonyl-CoA was added to each well, while the same quantity of molecular-grade water was added to background wells. FAS activity was then measured as the oxidation of NADPH at 340 nm using a UV/VIS spectrophotometer (Biotek HT1) for 10 minutes at 37°C. A background correction was made for the oxidation of NADPH in the absence of malonyl-CoA.

Lipase activity was measured in thawed aliquots of the fat body supernatant using a fluorometric Lipase Activity Assay Kit (Cayman Chemical) as has been previously done with honey bee fat body tissue (Corby-Harris, Snyder and Meador, 2019). Finally, the total amount of soluble protein in each sample was measured with a BCA Assay Kit (Thermo Scientific) according to manufacturer’s instructions, primarily as a normalization factor. Both FAS and lipase activities were calculated as whole abdominal values (units of nmol/min/abdomen) and normalized to total protein in each sample (units of nmol/min/mg).

### Hypopharyngeal gland (HG) acini size quantification

The HGs of bees that were fed or not fed protein and lipid for 8 days in cages were measured to determine whether HG size differed depending on diet. Bees were collected from each cage, their heads removed, flash frozen in liquid nitrogen, and maintained at -80°C until their glands were measured. For each bee, the HGs were dissected into phosphate buffered saline (7 mM NaCl, 2.7 mM KCl, and 10 mM PO4, pH 7.4), stained using Giemsa stain for 7 minutes, and then photographed using a Leica M205C stereoscope with a Leica DFC450 camera using Leica Applications Suite v4.5. The area of 10 acini per bee was measured using ImageJ and the mean value per individual used for statistical analysis. The selection criteria used for acini in each image was that had clear margins indicating that they were in focus, had an attachment point visible to the collecting duct, and appeared to be of average size relative to other acini in the photograph.

Because the measurements were carried out by a researcher who was not blind to treatment identity, assessment of possible selection bias was carried out by randomly selecting 20 photos from the pool of 106 using the R package “random” that generates random numbers from atmospheric noise, giving those 20 images to an observer blind to treatment identity, and asking them to select acini in the photos based on the described selection criteria. Selected acini were then measured, and the measurements compared to those made by the researcher. Measured acini size did not significantly differ based on whether the acini were selected by the researcher or an observer blind to study conditions (Mann–Whitney *U* = 2854, *n*_1_ = *n*_2_ = 75, *P* = 0.8775).

### Abdominal lipid quantification

The abdominal fat body lipid content of 3, 8, 13, and 21-day-old bees and of bees fed or not fed protein was measured. For the age-based colony samples, bees were collected, flash frozen in liquid nitrogen, and maintained at −80°C until their fat bodies were dissected as described above, following a previous approach (Corby-Harris, Snyder and Meador, 2019). The tissue samples were Folch extracted overnight at 4°C in 2 mL of 2:1 chloroform:methanol with 25 mg/mL of butylated hydroxytoluene (BHT) to inhibit lipase-based lipid degradation and lipid oxidation. The samples were then briefly vortexed and centrifuged for 15 min at 4°C. The bottom chloroform phase was retained and dried using a CentriVap at room temperature. The dried chloroform-soluble fraction was subjected to a sulfuric acid-vanillin-phosphoric acid assay as described previously (Van Handel, 1985). Sample absorbances were evaluated against a standard curve of corn oil. For the diet experiment, methods were identical except that fat bodies were dissected prior to flash freezing in liquid nitrogen.

### Statistical analyses

To address the effect of age and replicate on abdominal FAS activity, normalized FAS activity, and normalized lipase activity, we used an ANOVA with age, replicate, and their interactions as factors. Since abdominal protein and abdominal lipase activity did not meet the assumption of normality even when log transformed, a Kruskal-Wallis test was used to assess the effect of age, while a Wilcoxon signed rank test was used for replicate effects.

Similarly, ANOVA was used for the effect of age class, feeding status, replicate, and their interactions on abdominal FAS activity. Since abdominal protein quantity did not meet the assumptions of normality, a Kruskal-Wallis test was used to test the effect of treatment group (fed nurse, starved nurse, fed forager, starved forager) and replicate with a post-hoc Dunn’s test with a Bonferroni correction for multiple comparisons for the family-wise error rate (FWER) used to determine which groups differed significantly from each other.

All analyses were carried out using Rstudio version 2023.06.1.

## Results

### Fat body lipogenic capacity declines with age

To look for evidence that decreased synthesis or increased breakdown of lipids in the fat body is responsible for lipid loss during the nurse-forager transition, we first collected age-marked bees from hives and isolated their fat body tissue to measure enzyme activity and protein levels. Abdominal FAS activity was significantly affected by both age (*F*_2,42_ = 10.984, *P* = 0.000177) and replicate (*F*_1,42_ = 10.636, *P* = 0.002205). Bees at 13 days old had significantly lower FAS activity than those at 3 (*P* = 0.0165921) and 8 days old (*P* = 0.4529725), while 3 and 8-day-olds did not differ significantly (*P* = 0.4529725).

As a normalization factor, we also measured total protein quantity in the same set of abdominal carcass samples (Fig. 1A). Abdominal protein quantity was not affected by age (Kruskal-Wallis test, Chi-squared =2.99, *P* = 0.22, df = 2) or replicate (Wilcoxon signed rank test, *P* = 0.8143). Since there was no significant trend, we normalized FAS activity to the amount of protein in each sample, producing a similar result. There was a significant effect of both age (*F*_2,42_ = 8.068, *P* = 0.00108) and replicate (*F*_1,42_ = 4.104, *P =* 0.04916) on FAS activity (Fig. 1). Bees at 13 days old had significantly lower FAS activity than those at 3 (*P* = 0.0051925) and 8 days old (*P* = 0.0030271), while 3 and 8-day-olds did not differ significantly from each other (*P* = 0.9804045).

**Fig. 1.**
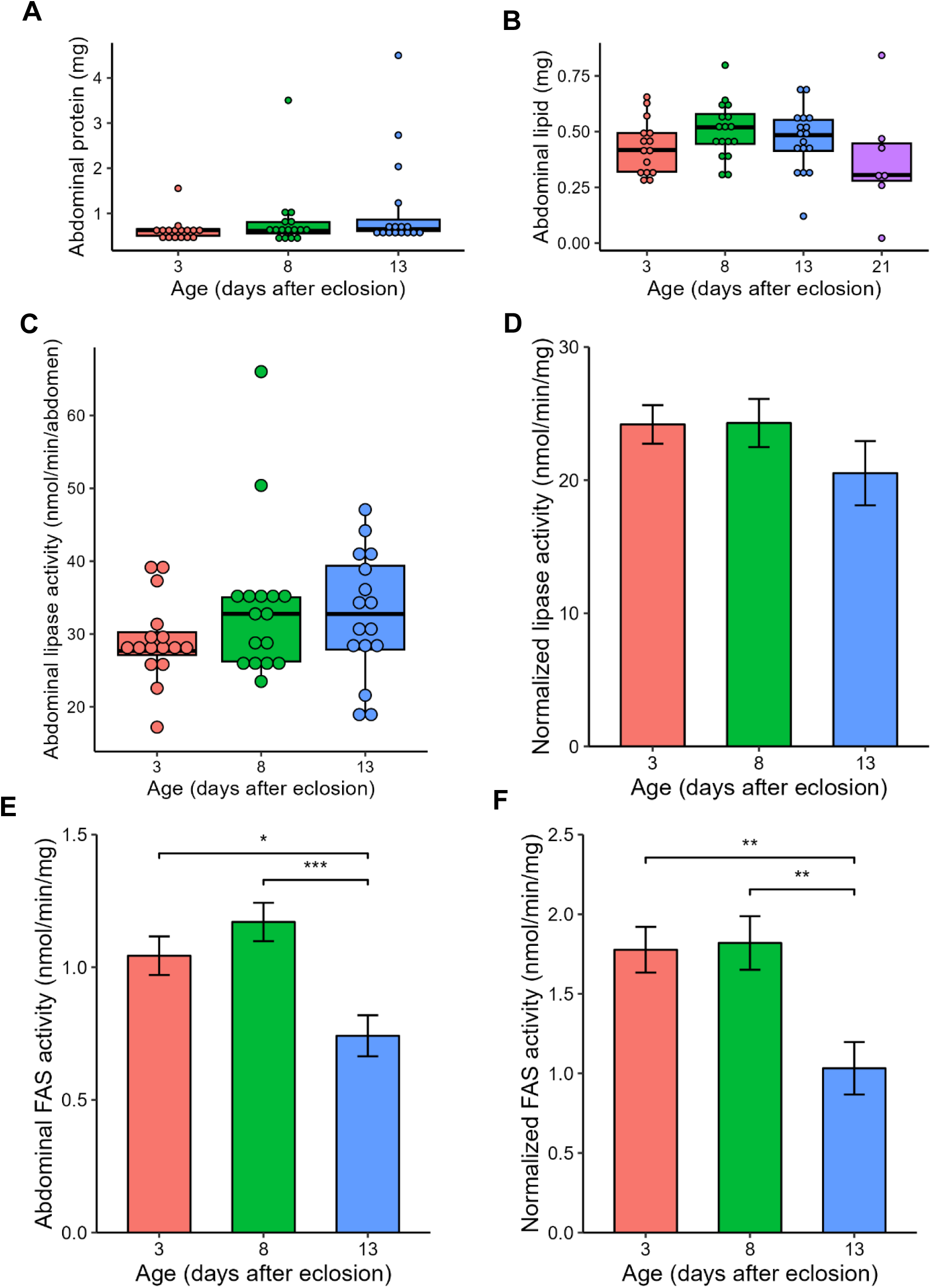
Lipogenic capacity declines as bees age past peak nursing age. (A) Abdominal protein does not change significantly with age. (B) Abdominal lipid quantity does not change significantly with age. (C) Abdominal lipase activity and normalized lipase activity do not change significantly with age. Activity levels are for C are in nmol of thioglycerol produced per minute per well while for D they are normalized to abdominal protein. (E) Abdominal fatty acid synthase (FAS) activity (F) normalized FAS activity are significantly lower in 13 day old bees compared to younger age groups. Activity levels for E are in nmol of NADPH oxidized per minute and for F are in nmol of NADPH oxidized per minute per mg of abdominal protein. Data that did not meet the assumption of normality are shown with dots as individual data points, and boxes, bars, and whiskers showing the upper and lower quartile range, median value, and minimum and maximum, respectively. For the data based on two pooled abdomens (A and C-F), *N=*16 samples of two pooled abdomens for each age group. For B, *N*=18, 20, 20, and 10, for 3, 8, 13, and 21-day-old bees, respectively. For normal data, bars represent the mean of each pooled age group ± SEM.

Abdominal lipase activity was not affected by age (Kruskal-Wallis, Chi-squared = 2.683, *P* = 0.2614, df = 2) or replicate (Wilcoxon signed rank test, *W* = 289, *P* = 0.9919; Fig. 1C). Similarly, normalized lipase activity was not significantly affected by age (*F*_2,44_ = 1.235, *P* = 0.30) or replicate (*F*_1,44_ = 1.434, *P* = 0.238; Fig. 1D). Abdominal lipids were not significantly affected by age (*F*_3,60_ = 1.353, *P* = 0.266) or replicate (*F*_1,60_ = 1.001, *P* = 0.321; Fig. 1B).

### Nurses have much higher lipogenic capacity than foragers and age class is more important than feeding status in determining lipogenic capacity

To test whether the lower FAS activity in 13-day old bees was due to a transition to foraging behavior or a difference in feeding status, we designed a two-factor experiment where 8-day-old nurses and 28-day-old foragers were either fed 30% sucrose solution *ad libitum* or starved (given only water) for 21 hours in cages. Abdominal FAS activity was significantly higher in nurses compared to foragers (*F*_1,44_ = 74.167, *P* < 0.0001) and in fed compared to starved bees (*F*_1,44_ = 8.556, *P* = 0.0054; Fig. 2B). Abdominal protein quantity was significantly lower in starved foragers than in the other three groups (Fig. 2A; Kruskal-Wallis, Chi-squared = 21.124, *P* < 0.0001, df = 3; post-hoc Dunn’s test with a Bonferroni correction for multiple comparisons for the family-wise error rate) and not significantly affected by replicate (Kruskal-Wallis, Chi-squared = 3.8903, *P* = 0.2736, df = 3) .Because abdominal protein quantity was significantly different across the groups, it was thus not used as a normalization factor and normalized FAS activity was not analyzed for this experiment.

**Fig. 2.**
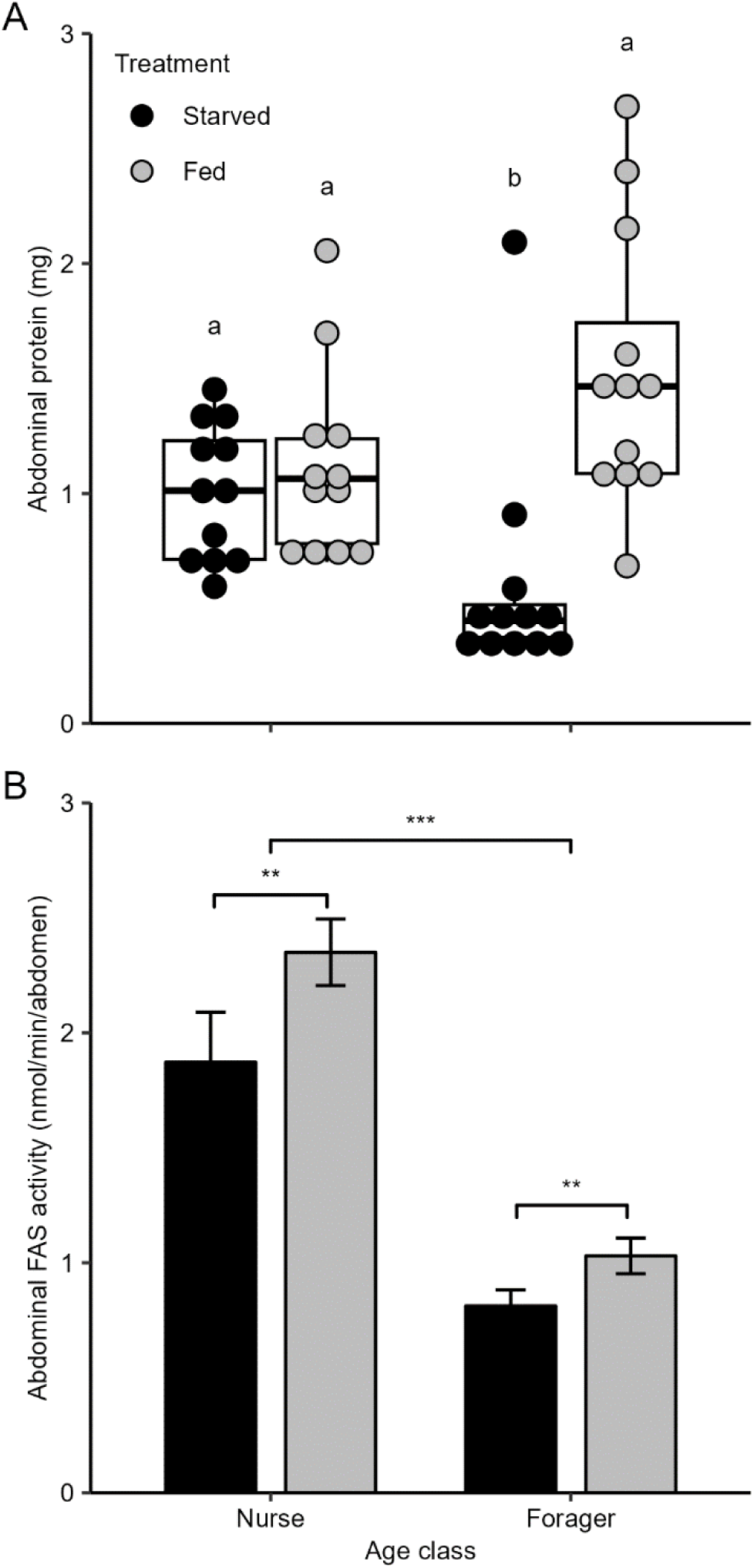
Both age-class and feeding status affect abdominal lipogenic capacity. (A) Starved foragers have significantly lower total abdominal protein than fed foragers and fed or starved nurses. Letters indicate the result of a post-hoc Dunn’s test with a Bonferroni correction for multiple comparisons for the family-wise error rate (FWER). *N*=12. (B) Abdominal FAS activity was significantly higher in nurses than foragers and in fed than starved bees. ANOVA, *N*=12, \*\**P≤*0.01, \*\*\**P≤*0.001. Bars represent the mean of each group ± SEM.

### Honey bees fed protein have higher fat body lipogenic capacity, more abdominal lipid, and larger hypopharyngeal glands

To test whether dietary composition could produce the differences in FAS activity between nurses and foragers, we designed artificial diets that allowed us to manipulate dietary protein and lipid content and fed these diets to bees in cages for 8 days. Total estimated calories consumed per bee per day was not affected by dietary protein (*F*_1,8_ = 2.415, *P* = 0.159) but was significantly higher in bees fed lipids (*F*_1,8_ = 5.436, *P* = 0.048; Fig. S1).

Abdominal protein was not significantly affected by dietary protein (*F*_1,20_ = 0.253, *P* = 0.621) nor dietary lipid (*F*_1,20_ = 2.119, *P* = 0.161; Fig. 3A). Abdominal FAS activity was significantly higher in bees fed protein compared to those deprived of protein (*F*_1,21_ = 63.781, *P* < 0.0001) but not significantly affected by dietary lipid (*F*_1,21_ = 0.795, *P* = 0.383; Fig. 3C). ANOVA of the log-transformed data showed that there was a significant effect of dietary protein (*F*_1,20_ = 53.752, *P* < 0.0001) and lipid (*F*_1,20_ = 6.530, *P* = 0.0189) on normalized FAS activity (Fig. 4C).

**Fig. 3.**
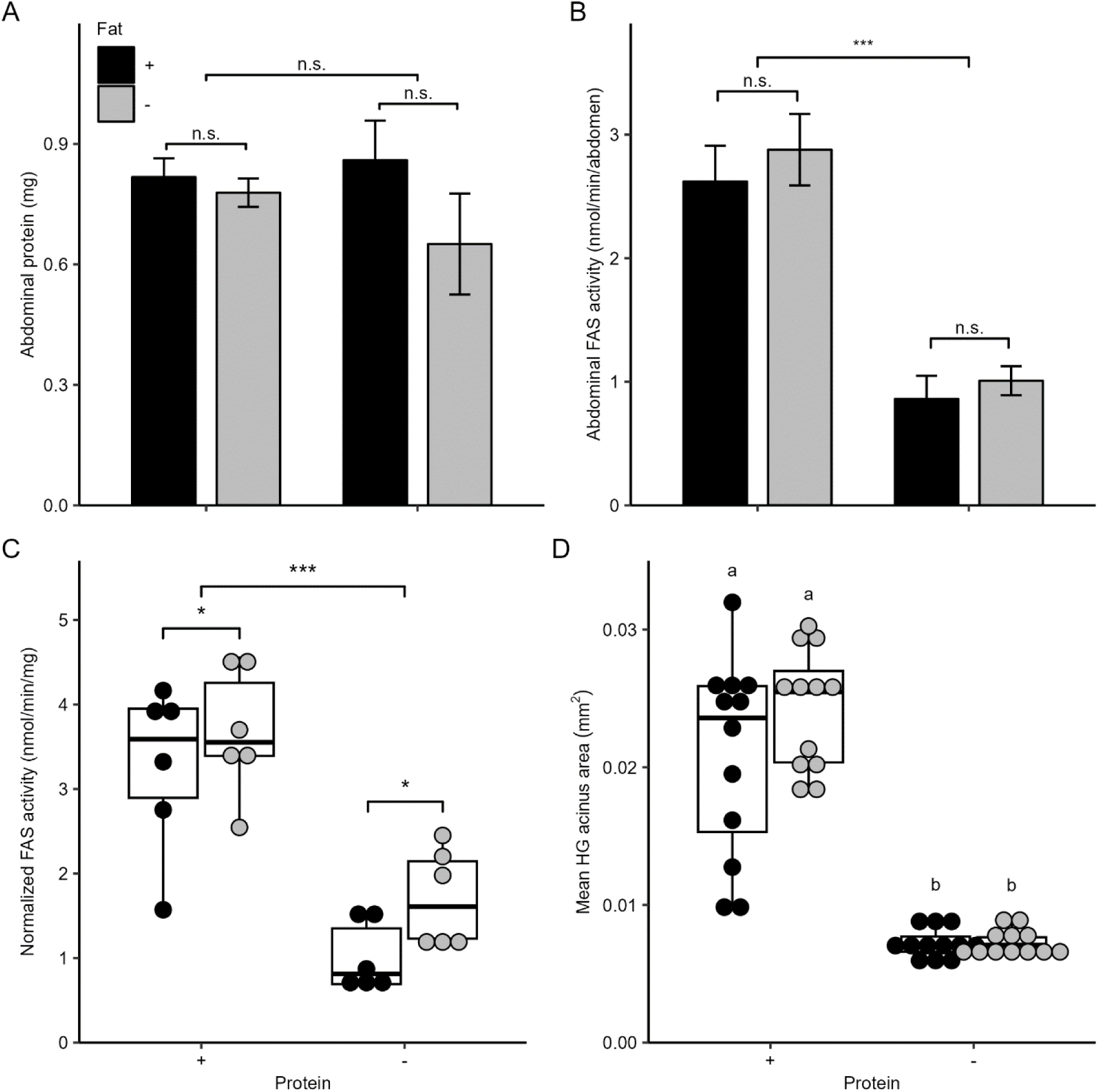
Dietary protein increases abdominal lipogenic capacity. (A) Dietary lipid and protein do not significantly affect total abdominal protein. *N*=6. (C) Abdominal FAS was significantly higher in bees fed protein but not affected by dietary lipid. ANOVA, *N*=6, \*\*\**P≤*0.001. (C) Normalized FAS activity was significantly increased by both dietary protein and fat. ANOVA, *N*=6, \**P≤*0.05, \*\*\**P≤*0.001. (D) Hypopharyngeal gland acini size is significantly larger in bees fed protein but not affected by dietary lipid. Letters indicate the result of a post-hoc Dunn’s test with a Bonferroni correction for multiple comparisons for the family-wise error rate (FWER). *N*=12. Data that did not meet the assumption of normality are shown with dots as individual data points, and boxes, bars, and whiskers showing the upper and lower quartile range, median value, and minimum and maximum, respectively. For normal data, bars represent the mean of each pooled age group ± SEM.

**Fig. 4.**
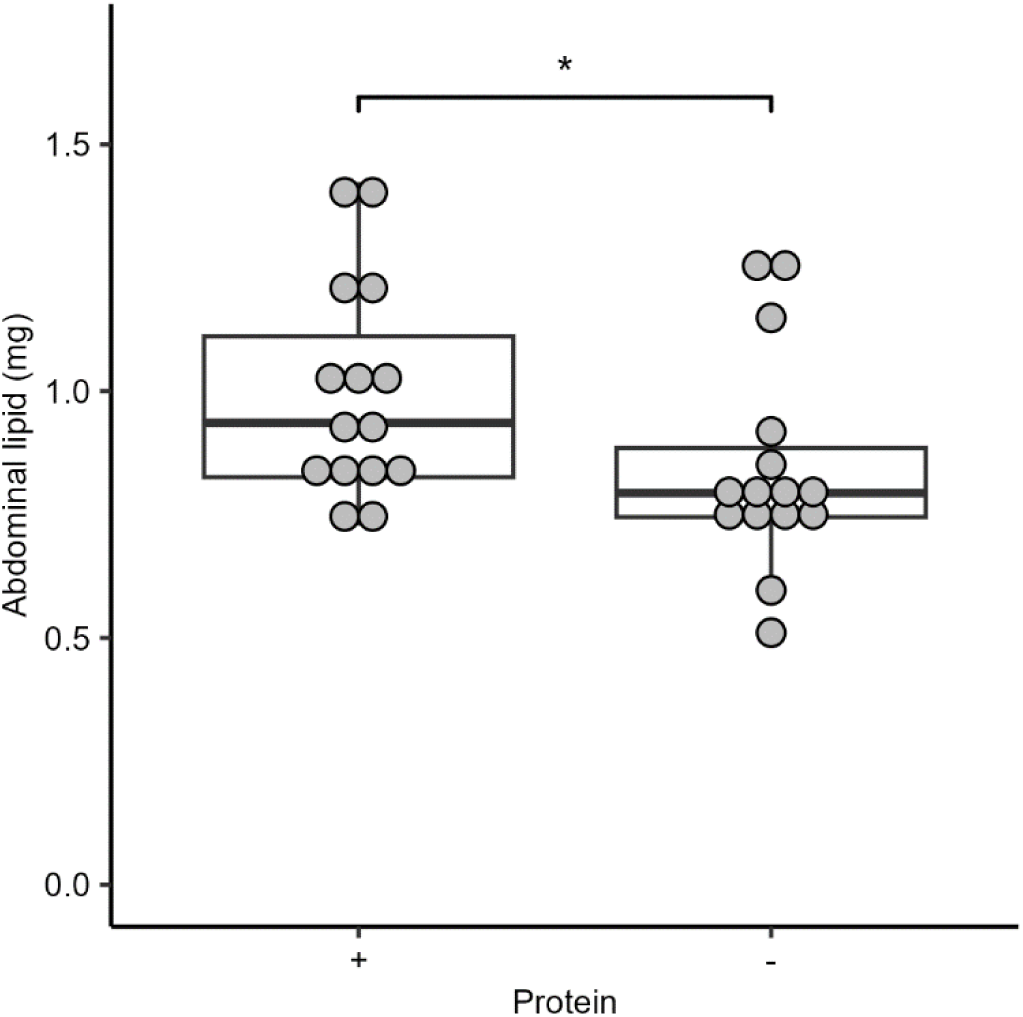
Abdominal fat is increased by a diet containing protein. When bees fed for 8 days with diets containing or lacking protein, bees fed protein have significantly higher total abdominal lipids. *N*=15, \**P≤*0.05 with Wilcoxon signed rank test.

To further address whether the diets produce bees with nurse-like physiology, we also measured the size of HGs and the total quantity of abdominal lipids. A Kruskal-Wallis test showed a significant difference in hypopharyngeal (HG) gland mean acinus area between treatment groups (Chi-squared = 35.586, *P* < 0.0001, df = 3; Fig. 3B). A post-hoc Dunn’s test with a Bonferroni correction for multiple comparisons for the family-wise error rate (FWER) showed that dietary protein but not fat increased HG gland mean acinus area. HG mean acini area was significantly higher in bees fed protein compared to those not fed protein and not significantly affected by dietary lipid (Fig. 3A; Kruskal-Wallis, Chi-squared = 21.124, *P* < 0.0001, df = 3; post-hoc Dunn’s test with a Bonferroni correction for multiple comparisons for the family-wise error rate). Abdominal lipid was significantly affected by replicate (Kruskal-Wallis, Chi-squared = 8.1626, *P* = 0.01689, df = 2). However, because the pattern across replicates was the same, data were pooled to determine treatment effects: protein-fed bees had significantly higher abdominal lipids than protein-deprived bees (Fig. 4; Wilcoxon signed rank test, *W* = 60, *P* = 0.0295).

## Discussion

In this study, we found that honey bee worker fat body lipogenic capacity was significantly affected by age, role within the colony, and dietary macronutrient availability. Lipogenic capacity declines as workers age past peak nursing behavior (8 days old; Fig. 1) and that, whether they are starved or fed, nurses have consistently higher lipogenic capacity than foragers (Fig. 2B). We also tested how availability of dietary macronutrients affects lipogenic capacity: it was significantly decreased by starvation in both nurses and foragers collected from colonies (Fig 2B) and by protein (but not lipid) deprivation for bees kept in a controlled environment in cages for 8 days (Fig 3B-C). Protein deprivation similarly decreased the size of HGs and total abdominal lipid content (Fig. 3A, D), matching past studies showing that sufficient protein consumption during development is necessary for workers to develop nurse-like physiology, while protein deprivation causes premature development of forager-like physiology (DeGrandi-Hoffman *et al*., 2010; Corby-Harris *et al*., 2016; Omar *et al*., 2017; Martelli *et al*., 2022).

Our finding here, that FAS activity and not lipase activity or total protein is downregulated in older workers and foragers, supports the hypothesis that reduced capacity to synthesize lipids is specifically targeted for downregulation during the nurse-forager transition. Reduced lipogenic capacity provides one mechanistic explanation for why foragers become and remain lean, despite consistently maintaining higher hemolymph sugar levels than nurses (Mayack *et al*., 2019). A limitation of our study is that, because we used a spectrophotometric assay for measuring the activity of the FAS enzyme complex rather than directly measuring the rate of lipogenesis itself, we cannot conclude definitively that stored fat body TAG correlates with the measured FAS activity. Our measurement of total abdominal lipid content provides some support for this correlation, since treatment groups with higher FAS activity generally had higher total abdominal lipids (Figs. 1B and 4). However, it should be noted that we did not conclusively show that FAS-derived fatty acids are stored as TAG in fat body lipid droplets: synthesized fatty acids may instead be directly catabolized via beta oxidation, used as a source for biosynthesis of lipid molecules used for cuticular hydrocarbons, tracheal waterproofing, cell membranes, or hormones (Huang, Liu and Perrimon, 2022), or transported by lipoproteins to the head glands to be used for the fatty acid fraction of jelly fed to developing bees.

The differences we found in lipogenic capacity between nurses and foragers persist even when carbohydrate feeding status is controlled by feeding bees *ad libitum* for 24 hours in a controlled laboratory environment: while acute starvation did decrease abdominal FAS activity in both nurses and foragers, age class had a much larger effect. Feeding on carbohydrates thus clearly upregulates lipogenic capacity, but it does so within a dynamic range determined by the social role of the bee. Interestingly, while in nurses the downregulation of FAS activity during starvation seems to be specific, with no reduction in total abdominal protein, starved foragers had significantly reduced total protein in the abdominal fat body relative to fed foragers and to nurses (Fig. 2). This suggests that the downregulation of FAS activity in starved foragers is less specific and part of a broad metabolic response to catabolize fat body proteins to provide energy during starvation. The lack of this response in nurses could be because they lack the necessary proteolytic machinery necessary for rapid catabolism of protein reserves, or because they are better buffered against nutritional stress, perhaps because of their higher lipid stores (Toth and Robinson, 2005), higher levels of hemolymph storage proteins such Vitellogenin that can be catabolized instead of fat body protein (Nakaoka, Takeuchi and Kubo, 2008), or their lower maximal metabolic rate (Schippers, Dukas and McClelland, 2010) or lower activity level. Further work will be needed to address age-class specific differences in protein catabolism.

Our finding that protein deprivation causes reduced lipogenic capacity (Fig. 3C-D), lower abdominal lipids (Fig. 4), and smaller HGs (Fig. 3B) in nurse-age bees fits into a broad range of research linking dietary protein with development and maintenance of nurse physiology. A nurse’s function in the colony is primarily to produce and distribute royal jelly to the queen, larvae, and other workers, acting as a net nutritional source while other bees are nutritional sinks (Crailsheim, 1991). To support synthesis of royal jelly, nurses are the primary consumers of pollen in a hive (Crailsheim *et al*., 1992) and they have much higher levels of digestive proteases than other bees so that they can process difficult to digest pollen into useful nutrition (Crailsheim and Stolberg, 1989). Royal jelly is a complete food source composed of protein, carbohydrates, and lipids (Wang *et al*., 2016). It is secreted from the HGs and mandibular glands (MGs) of the head. Adequate dietary protein during the first week of life is critical for the development of the physiological features enabling effective nursing behavior: in bees deprived of pollen, the HGs degrade between 3 and 8 days (Corby-Harris *et al*., 2022), and our replication here of this pattern using artificial diets confirms the protein component of pollen as the primary factor necessary for the development of large, nurse-like HGs. Pollen deprivation similarly decreases total HG and MG protein (Peters, Zhu-Salzman and Pankiw, 2010) and Vitellogenin hemolymph titer (Bitondi and Simões, 1996) and there is a positive correlation between pollen in the worker intestines and fat body Vitellogenin (Wegener *et al*., 2018). Vitellogenin is used by nurse bees as an amino acid donor for producing royal jelly and high Vitellogenin titers are linked with nursing (Amdam *et al*., 2003). Since successful nursing behavior requires protein, it is beneficial for colonies if nurses that do not have sufficient dietary protein transition to foraging to increase pollen and hence protein flow into the hive (Amdam and Page, 2005).

While here we found no effect of dietary lipids on HG size (Fig. 3B), another study found that feeding bees lipids did increase HG size (Stabler *et al*., 2020). The difference may be due to the older age of the bees in that study (10 days), the different lipid source (soy lecithin instead of corn oil), or differences in consumption based on the state of the diets (liquid instead of semi-solid). Lipids are an important but variable part of royal jelly, ranging from approximately 2-7% in one study (Ferioli, Armaforte and Caboni, 2014). This variability is likely because lipids are the most variable composition in pollen, ranging from 1-13% (Campos *et al*., 2008). Lipids are likely a valuable resource to colonies, since increasing the dietary lipid concentration causes nurse-age bees to consume more total food (Stabler *et al*., 2020) and have increased success in brood-rearing (Arien *et al*., 2020).

We found mixed evidence regarding whether lipogenic capacity is affected by dietary fat. While groups fed lipids generally had lower FAS activity, this difference was only significant when normalizing to total protein (Fig. 3B-C). Caution should be taken in interpreting this result, since, while the difference was not significant, bees deprived of lipids had lower total abdominal protein, potentially biasing the result (Fig. 3A). It is worth nothing that bees fed lipids consumed more total calories (Fig. S1), a finding that is consistent with past work (Stabler *et al*., 2020), suggesting that any potential upregulation in FAS activity in bees deprived of lipids is not due to increased consumption. It is possible that workers increase *de novo* lipogenesis to compensate for deficiency in dietary lipids or decrease lipogenesis when dietary lipids are adequate, but if so, the effect is very small compared to the effect of dietary protein.

Vitellogenesis has been evolutionarily co-opted in honey bees to provide larval nutrition from a sterile worker class (Amdam *et al*., 2003). Much of nurse physiology seems to be optimized for high levels of fat body vitellogenesis, and this may extend to the high nurse fat body FAS activity we measured here as well: in planthoppers, knockdown of FAS lowered expression of both *Vg* and *VgR* (Cheng *et al*., 2023), suggesting that *de novo* synthesized fatty acids are used for vitellogenesis, possibly because Vitellogenin requires lipidation by specific fatty acids before secretion (Silversand and Haux, 1995).

Our finding that dietary protein deprivation prematurely induces a forager-like reduction in lipogenic capacity and fat body lipids suggests that the dietary switch during the nurse-forager transition could be an important upstream signal lowering synthesis and storage of lipids in the fat body. This study does not directly address why foragers consume less protein that nurses, but a range of studies support that that they do: experiments using paired diets in a geometric framework that showed that the intake target of young, queenless bees shifted from a proportion of EAAs (essential amino acids)-to-carbohydrates (EAA:C) of 1:50 towards 1:75 over a 2-week period, accompanied by a reduced lifespan on diets high in EAAs. Foragers required a diet high in carbohydrates (1:250) and also had low survival on diets high in EAA (Stabler *et al*., 2020). Foragers consume little or no pollen (Crailsheim *et al*., 1992), and thus jelly received from nurses must be their primary source of protein (Crailsheim, 1992). The amount of jelly given to adult workers does significantly decrease with age, suggesting that foragers receive less total protein than younger workers (Crailsheim, 1992).

While we focused on synthesized lipids in this study, dietary lipids also likely play a role, since foragers consume less pollen than nurses, and pollen is the primary source of dietary lipids. We did not address the relative contributions of dietary and *de novo* lipids to total lipid stores in workers, but past work has shown that treating workers with a fatty acid synthesis inhibitor significantly decreases abdominal lipids only when workers are deprived of pollen (Toth *et al*., 2005), suggesting that *de novo* lipogenesis becomes more important for lipid storage when dietary lipids are not available. Since foragers consume minimal amounts of pollen (Crailsheim *et al*., 1992), reduced lipogenic capacity is likely especially important in maintaining lipid stores at the low levels found in foragers.

Taken together, our findings implicate a developmental reduction in protein intake targets as a central factor in mediating metabolic plasticity in honey bee workers. This support the hypothesis that honey bee worker division of labor evolved through co-option of, and novel, social inputs into, the regulation of appetite, but much more experimental work will be needed before the synthesis of a complete model linking hive inputs through regulation of protein appetite into mediation of caste-specific metabolic physiology.

## Acknowledgements

We thank Cahit Ozturk for his assistance with bee husbandry, Jenna Dobson for help with HG acini quantification, Vanessa Corby-Harris for advice on lipase activity and lipid quantification assays, and Geraldine Wright and Daniel Stabler for help with the design of artificial diets. We thank Tim Linksvayer for his helpful comments on the manuscript.

## Data availability

Data are temporarily available via this private Figshare link: https://figshare.com/s/126451f85e548b4f0208

Data will be published and made publicly accessible via Figshare at the time of publication.

